# Deciphering structural complexity of brain, joint, and muscle tissues using Fourier Ptychographic Scattered Light Microscopy

**DOI:** 10.1101/2024.11.28.625428

**Authors:** Simon E. van Staalduine, Vittorio Bianco, Pietro Ferraro, Miriam Menzel

## Abstract

Fourier Ptychographic Microscopy (FPM) provides high-resolution imaging and morphological information over large fields of view, while Computational Scattered Light Imaging (ComSLI) excels at mapping interwoven fiber organization in unstained tissue sections. This study introduces Fourier Ptychographic Scattered Light Microscopy (FP-SLM), a new multi-modal approach that combines FPM and ComSLI analyses to create both high-resolution phase-contrast images and fiber orientation maps from a single dataset. The method is demonstrated on a state-of-the-art setup that was originally used for FPM and one that was originally used for ComSLI, and the outputs are quantitatively compared to each other on brain sections (frog, monkey) and sections from thigh muscle and knee (mouse). FP-SLM delivers high-resolution images while revealing fiber organization in nerve, muscle, tendon, cartilage, and bone tissues. The approach is validated by comparing the computed fiber orientations with those derived from structure tensor analysis of the high-resolution images. The comparison shows that FPM and ComSLI are compatible with each other and yield fully consistent results. Remarkably, this combination surpasses the sum of its parts, so that applying ComSLI analysis to FPM recordings and vice-versa outperforms both methods alone. FP-SLM can be retrospectively applied to analyze any existing dataset acquired from a setup that was originally built for FPM or ComSLI alone (consisting of LED array and low numerical aperture), without need to build or design an extra setup. This significantly expands the application range of both techniques and enhances the study of complex tissue architectures in biomedical research.

## 1 INTRODUCTION

In biomedical research, both broad tissue context and cellular-level resolution are essential to analyze tissue morphology and pathology. Super-resolution methods like stimulated emission depletion (STED) achieve molecular-level detail [1, 2] but require fluorescent labeling, which may alter biological processes and requires complicated sample preparation, and they only provide a small field of view. Confocal or two-photon microscopy achieve sub-micrometer resolution [3, 4] but are also limited by slow acquisition speed and small field of view. High-resolution imaging requires high-magnification lenses, but these restrict the field of view, making it difficult to observe large tissue areas while maintaining fine details.

*Fourier Ptychographic Microscopy* (FPM) is a label-free technique that combines low-resolution intensity images captured under different illumination angles to reconstruct high-resolution complex amplitude images over large fields of view [5]. In this way, it overcomes the traditional trade-off between resolution and field of view in conventional microscopy, allowing for detailed imaging of biological tissues without the need for high magnification optics. FPM enables quantitative phase imaging of large histological sections with cellular-level detail, making it ideal for studying complex tissue samples like brain or muscle tissues [6, 7]. Accessing the phase map from intensity-only captures allows measuring biophysical parameters of the specimen, since the phase contrast is proportional to the optical thickness and dry mass. As any stain-free method, FPM lacks specificity since the contrast mechanism it relies on is endogenous. Indeed, the optical readout is the difference in the phase delay experienced by a light wavefront when passing through the sample with respect to its unperturbed propagation through the surrounding medium. The lack of specificity can be considered as the main shortcoming of quantitative phase imaging approaches and much effort has recently been spent to bridge the gap between stain-based and stain-free approaches using, e.g., artificial intelligence [8, 9, 10] or computational microscopy methods [11, 12].

While FPM provides phase-contrast in unstained tissues, it is not suited for studying the structural organization of tissue because differently oriented structures with similar optical thicknesses yield similar contrasts. Moreover, the optical thickness is not univocally associated with the presence of fiber bundles, so that areas with high optical thickness contrast in a tissue region can be ascribable to the presence of various elements with large refractive index, e.g. cells nuclei, lipid droplets, lysosomal accumulation, or fat areas. Yet, the orientation of structures such as nerve, muscle, or collagen fibers contains information about structure and pathology of the tissue that is essential for histological research and diagnostics. The brain, for example, consists of densely packed nerve fibers, and disentangling this complex fiber network – the human connectome [13] – is crucial for understanding the brain’s function in health and disease. Collagen fibers are abundant in the human body and can be found among others in tendon, cartilage, bone, or connective tissue. In cancerous tissues, the orientation of collagen fibers with respect to the tumor nest is related to tumor invasion and progression [14, 15, 16], determining the severity of cancer. For a detailed histological analysis, it is thus not only important to study tissue morphology at high resolution but also to determine the orientation of tissue structures.

To analyze the organization of nerve or muscle fibers, structure tensor analysis (STA) [17] can be applied to the high-resolution phase maps from FPM. However, it falls short in the case of dense tissues and regions where several fibers cross each other, and – being an indirect method – it can be prone to errors. In dense tissues, where the fibers are tightly packed, the intensity gradients used by STA can become ambiguous in FPM. In fact, integral phase signals in FPM from fibers located at different depths can blur or distort the local orientation information, reducing the accuracy of the analysis. A direct way to analyze fiber organization is polarization microscopy, which measures the optical anisotropy (birefringence) of the tissue to determine the orientation of the optic axis, corresponding to the fiber orientation [18]. Polarization microscopy has been established for decades, and is current gold standard in research and clinical laboratories for analyzing directed structures in histological tissue samples. However, the technique is limited to birefringent samples [19], and only yields one fiber orientation per image pixel [20], making it difficult to study the manifold interwoven fiber structures in biological tissues.

*Computational Scattered Light Imaging* (ComSLI) overcomes these limitations. By analyzing scattering patterns of light, it allows to accurately reconstruct multi-directional fibers, also in regions with densely packed, interwoven fibers like brain tissue [21, 22]. Unlike polarization microscopy, ComSLI reveals directed structures also in tissues with little or no birefringence, like paraffin-embedded brain tissues [19]. ComSLI is a label-free technique and works on both stained and unstained tissues with different tissue preparation protocols. It provides detailed fiber organization maps over large fields of view, achieving pixel sizes of less than 3 µm and fields of view of more than 1.5 cm^2^ [19], but it still has a lower resolution than FPM for the same field of view and does not provide optical density information. Hence, there is a need for a technique that provides both biophysical parameters with cell-level details and interwoven fiber organization in the same tissue sample.

**Figure 1** shows the principle of FPM and ComSLI. They both use the same basic setup and measurement principle (a): the sample is illuminated from an array of LEDs at different angles of incidence, and the transmitted light is recorded for each illumination angle by a camera using an optical system with small numerical aperture (NA), yielding a stack of images (b). FPM makes use of the fact that, for each illumination angle, a different part of the sample’s frequency spectrum is imaged (c1). The images are stitched together in Fourier space using a gradient-descent optimization algorithm like EPRY which also serves as an iterative phase-retrieval solver (c2), and transformed back into a high-resolution real-space image; the resulting image is complex-valued, of which only the phase is displayed here (phase map, c3). Instead of representing each image in frequency space, ComSLI analyzes the scattering patterns for each image pixel (d1): the data is rearranged so that each pixel in (*x, y*) space shows the intensity response per LED position in (*u, v*) space. As light scatters mostly perpendicular to directed structures like fibers, the peaks in the scattering pattern of each image pixel can be used to determine the local fiber orientations which are displayed in different colors (dashed lines in d1); multiple orientations per pixel are represented by a multi-colored pixel (d1,II). All pixels in the field of view are evaluated this way, resulting in a fiber orientation map (d2), enabling a detailed analysis of fiber organization in biological tissues, especially in regions with densely interwoven fibers.

**Figure 1:**
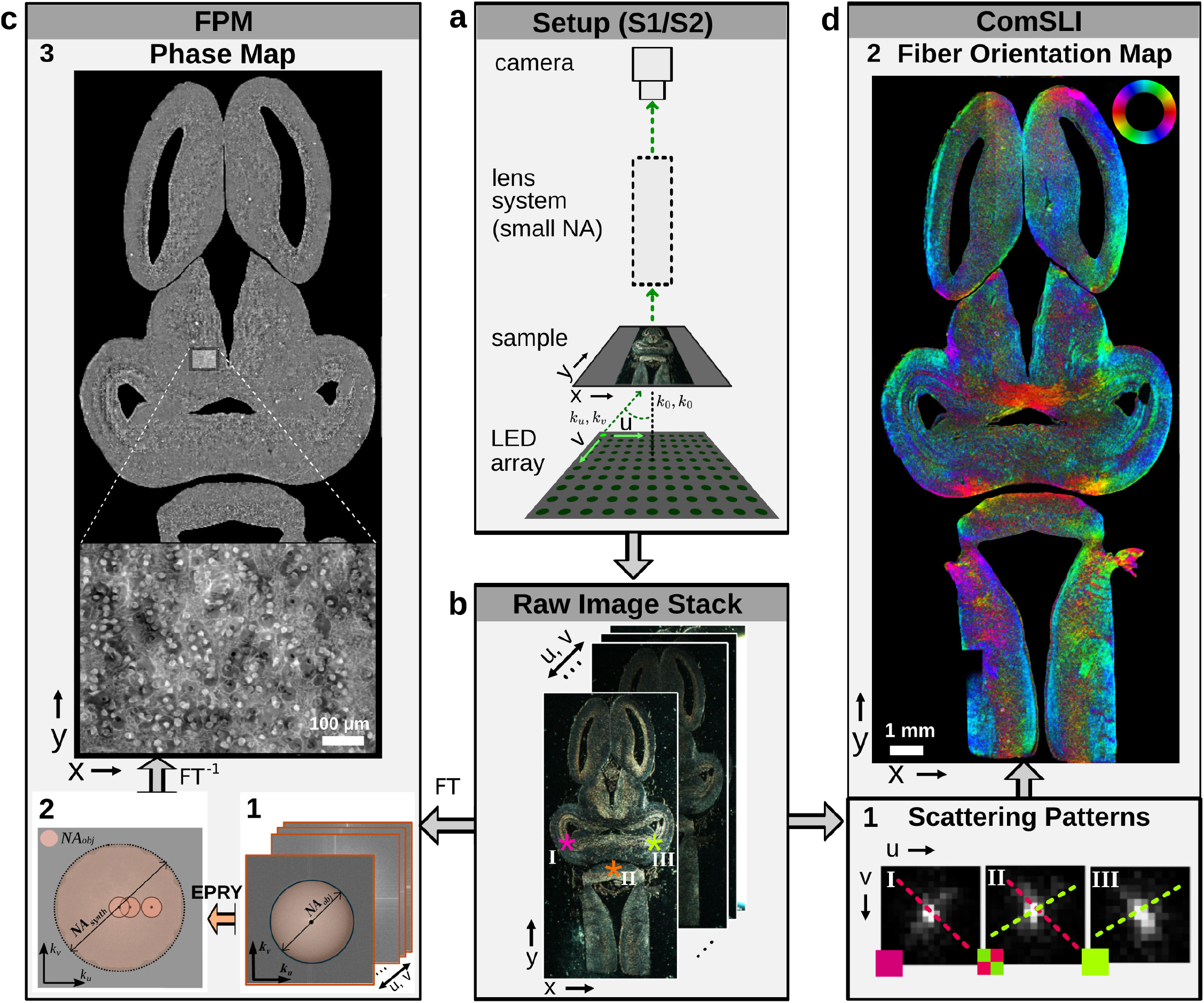
Principle of FPM and ComSLI, shown exemplary for a frog brain section. a) Both techniques use the same generic setup, consisting of an LED array that illuminates the sample under different angles and an optical system with small numerical aperture (NA). b) For each illumination angle (*u, v*), the camera records an image, yielding a stack of raw images. c) FPM applies a Fourier transform (FT) to the raw image stack (c1), uses a gradient-descent optimization algorithm like EPRY to stitch the images together in frequency space (c2), and transforms this image back into a high-resolution real-space image (phase map, c3). d) ComSLI rearranges the raw image stack to generate scattering patterns for each image pixel (d1) marked by colored asterisks in b; from the peak positions in the scattering pattern, the fiber orientations (dashed lines) are computed, which are then represented in different colors according to a color wheel – pixels with multiple fiber orientations are represented by a multi-colored pixel (d1,II). All colored pixels together yield the fiber orientation map (d2).

Since FPM and ComSLI use the same basic setup (LED array with low-NA objective), this offers the opportunity to combine the advantages of both methods. So far, FPM analysis has only been applied to data from setups built for FPM in order to generate high-resolution real-space images, and ComSLI analysis has only been applied to data from setups built for ComSLI in order to compute fiber orientation maps.

Here, we present the *Fourier Ptychographic Scattered Light Microscopy* (FP-SLM), a multi-modal approach that combines FPM and ComSLI analysis on the same dataset. We show that ComSLI analysis can be applied to data originally used for FPM to generate a fiber orientation map, and FPM analysis can be applied to data originally used for ComSLI to generate a high-resolution real-space image (phase map) in addition to the fiber orientation map provided by standard ComSLI analysis. We demonstrate the performance of FP-SLM on two state-of-the-art setups that have originally been designed for FPM and ComSLI, respectively (see Supplementary Figure 2).

Our results show the capability of FP-SLM to improve the performance of FPM and ComSLI alone. To demonstrate the potential for analyzing tissues across different types and species, we analyze brain sections from a frog tadpole and a vervet monkey, as well as skeletal muscle and knee joint sections from a mouse leg, containing different commonly studied tissue types: nerve, muscle, tendon, cartilage, and bone. For both setups, we compare the corresponding ComSLI fiber orientation maps, and the corresponding FPM high-resolution phase maps. We show that the ComSLI method for generating fiber orientation maps can also be employed on a setup originally used for FPM, and a setup originally used for ComSLI can also perform FPM, because both ComSLI and FPM access the same areas in frequency space. Thus, FP-SLM outperforms both FPM and ComSLI in terms of information channels one has access to. To study the performance of FP-SLM, a quantitative comparison is made between the ComSLI fiber orientations and the orientations retrieved from an STA analysis of the FPM phase maps.

## 2 RESULTS

Four different samples from three different species were analyzed: two samples were obtained from a mouse leg, cut longitudinally along the thigh bone, with one sample only containing skeletal muscle and the other sample also containing knee joint and bone. In addition, a horizontal section of a frog tadpole brain was studied which contains a great extent of gray matter, and – for comparison with STA – a coronal section of an adult vervet monkey brain which contains a lot of white matter fiber crossings in the corona radiata region. While the vervet brain section is a 60 µm-thin, unstained cryo-section used for previous ComSLI studies [22, 23], the mouse leg and frog brain samples are 4 µm- and 10 µm thin, respectively, formalin-fixed paraffin-embedded (FFPE), and stained with hematoxylin & eosin (H&E) and Golgi-stain, respectively, see Methods for more details. Supplementary Figure 1 shows the mouse knee and the frog and vervet brain samples together with some anatomical descriptions. The samples were measured with two state-of-the-art setups (see Supplementary Figure 2):

*Setup S1* has originally been designed for FPM [24, 25]. It is infinity corrected with a low-NA microscope objective (NA_obj_ = 0.1, 4.29 × magnification), and employs a circular LED array with 15-LED diameter which illuminates the sample under a maximum polar angle of 31° and generates a stack of 177 images with 2.05 × 1.55 mm^2^ field of view. The raw low-resolution intensity images (and the fiber orientation map obtained from ComSLI) have a pixel size in the image plane of 1.06 µm and a lateral optical resolution of at least 2.76 µm, corresponding to the low numerical aperture of the employed microscope objective. After the synthesis of all 177 images, the reconstructed high-resolution phase map has a pixel size in the image plane of 0.21 µm and a lateral optical resolution of at least 0.49 µm, corresponding to a synthetic NA_synth_ = 0.6 which allows to resolve fine structural details (see Supplementary Figure 2c). To measure a larger region, the sample was moved in the x- and y-directions, and the resulting phase maps were stitched together.

*Setup S2* has originally been designed for ComSLI [23]. It is not infinity corrected with a low-NA microscope objective (NA_obj_ = 0.045, 0.83 × magnification), and uses a square array of 40 × 40 LEDs illuminating the sample under a maximum polar angle of 20° and generating a stack of 1600 images with 16 × 11 mm^2^ field of view. The raw images and the corresponding ComSLI fiber orientation map have a pixel size in the image plane of 2.87 µm and a lateral optical resolution of at least 4.38 µm. After the FPM reconstruction of the raw images, the resulting phase map has a pixel size in the image plane of 0.56 µm and a lateral optical resolution of at least 0.87 µm (see Supplementary Figure 2f).

The samples were measured with both setups, S1 and S2. For each resulting dataset, the phase map and the fiber orientation map were generated using FPM and ComSLI analysis, respectively. Subsequently, STA was applied to the phase maps. The resulting maps were compared to each other qualitatively, and difference maps and histograms were computed for quantitative analysis, see Methods for more details. To generate fiber orientation maps from the scattering patterns (Figure 1d), a new filtering approach was used in which smoothed azimuthal line profiles were computed from the scattering patterns, enabling to more precisely determine the positions of high-intensity peaks and thus the local fiber orientations (see Methods and Supplementary Figure 4).

### 2.2 Reconstructed fiber orientations

First, the ComSLI fiber orientations obtained from both setups, S1 and S2, were compared to each other (see **Figure 2**). The orientations are shown in different colors according to the color wheel displayed in the subfigures (red corresponds to a horizontal orientation, cyan to a vertical orientation within the image plane). Different tissue sections were analyzed: frog tadpole brain (top, a), mouse thigh muscle (middle, e), and mouse knee (bottom, h). Apart from a side-by-side comparison (b,f,i), difference maps (d,j), and histograms for muscle, cartilage, tendon and bone (k) were computed for a quantitative analysis. The ComSLI fiber orientation maps obtained from setup S1 (originally designed for FPM) look strikingly similar to those obtained from setup S2 (originally designed for ComSLI). Observed differences are mostly due to the different resolutions: Setup S1 employs a 4.29 × magnification whereas setup S2 has a 0.83 × magnification. This becomes especially evident in the skeletal muscle sample (Figure 2g): while setup S1 resolves the substructure of skeletal muscle discs as horizontal stripes (orange arrows), setup S2 shows them as crossing substructures indicated by red-cyan subpixels. The knee joint sample contains different types of fibrous tissue (marked in Figure 2h) which can be used for direct comparison: muscle tissue containing muscle fibers (M), as well as cartilage (C), tendon (T), and bone (B) tissues containing mostly collagen fibers. All regions show a strong visual similarity in orientation colors. The muscle regions show again the horizontal stripes for setup S1 and the crossing substructure for setup S2 (see Figure 2i, regions I and III).

**Figure 2:**
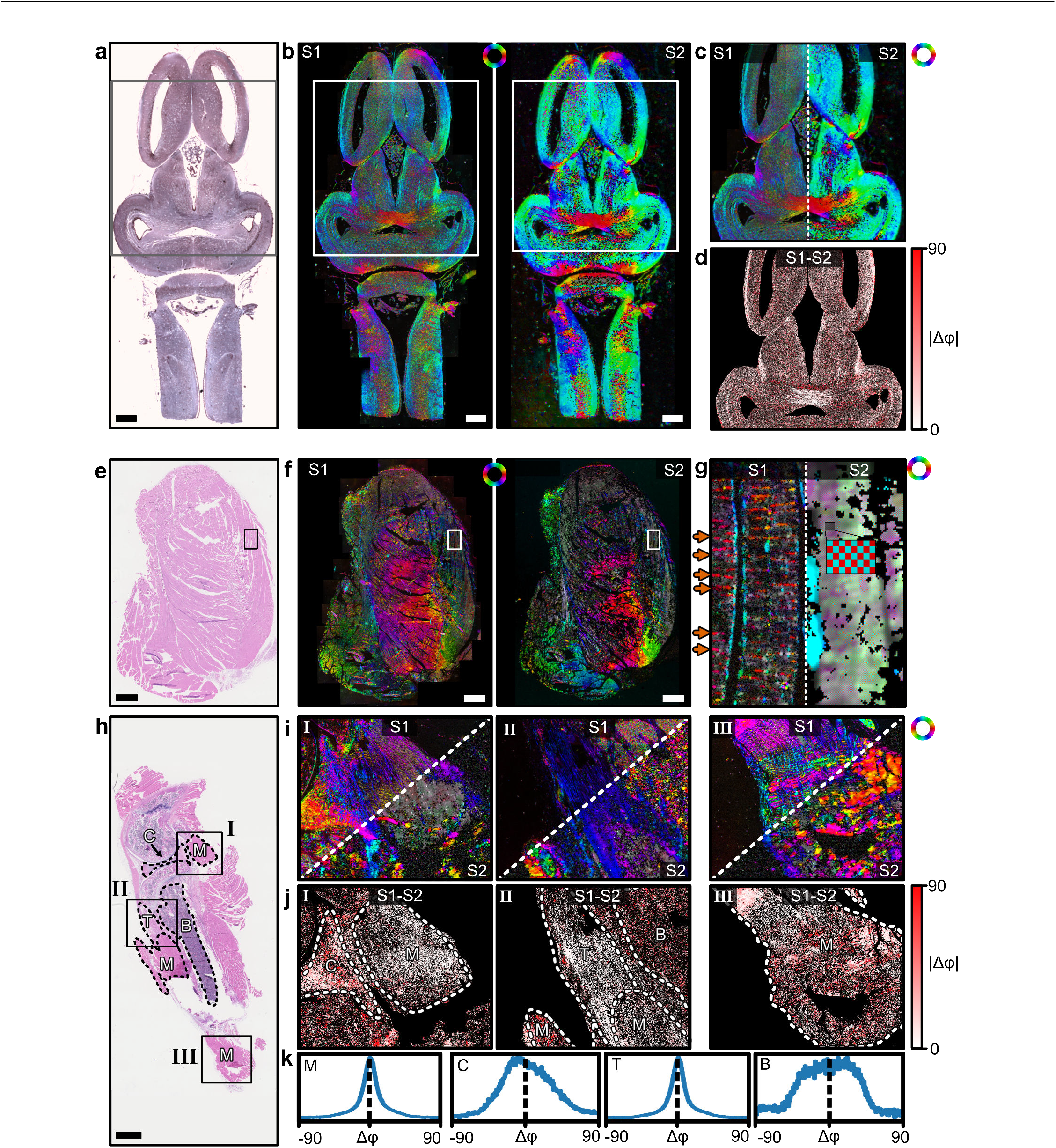
Comparison of ComSLI fiber orientation maps obtained from setup S1 (originally designed for FPM) and setup S2 (originally designed for ComSLI), shown for different tissue sections: frog brain (top), mouse thigh muscle (middle), and mouse knee (bottom). a),e),h) Bright-field microscopy images of the respective histology sections. M–muscle, C–cartilage, T–tendon, B–bone. b),f) Comparison between ComSLI fiber orientation maps from setup S1 (left) and setup S2 (right) for the brain and thigh muscle sample. c),g),i) Split-views of the corresponding fiber orientation maps for the rectangular regions marked in the histology sections. The orange arrows in (g) point towards horizontal (red) stripes which are related to the disc-structure in the skeletal muscle which is resolved by setup S1; the zoomed-in checkerboard pattern shows dual-colored pixels in this region for setup S2, indicating two perpendicularly running structures. d),j) Absolute difference between the fiber orientations obtained from both setups for the regions in (c) and (i), computed for each image pixel in degrees. k) Histograms of difference values evaluated separately for each tissue type (M–muscle, C–cartilage, T–tendon, B–bone; see regions marked in j). Histogram metrics (mean, mode, full width at half maximum, root mean square error) are listed in Supplementary Table 1. Scale bars correspond to 1 mm.

Quantitative similarity between measurement datasets is shown in the absolute difference maps below the different regions of interest (Figure 2d,j). In the frog brain sample, it can be seen that the differences in computed fiber orientations depend on the region: in regions with nerve fiber bundles (like in the bottom middle), there is much better agreement between setup S1 and S2 than in the surrounding gray matter regions with less aligned structures and lower scattering signal. In the knee sample, differences between tissue types become evident: while regions with cartilage (I, left) and bone (II, upper right) show larger absolute differences between the computed fiber orientation values, regions with muscle and tendon show smaller absolute differences. When comparing the histograms of the different tissue types (Figure 2k), the regions with muscle fibers and tendon (collagen fibers) yield histograms with a clear peak around zero and a full width at half maximum of about 18° (see Supplementary Table 1 for the complete metrics). For all fiber types – nerve fibers in the frog brain as well as muscle and collagen fibers in the mouse leg samples – there is a clear correspondence between the fiber orientations reconstructed from both setups. This is a first important result of our analysis, which shows the substantial compatibility between the setups that were originally used for FPM and ComSLI analysis alone. Cartilage and especially bone have a much less oriented substructure than muscle or tendon (as becomes visible in the phase maps shown later in Figure 3b). In the comparison of the two setups, we see an overall resemblance in the ComSLI fiber orientation maps, while the pixel-wise difference maps and histograms show larger differences for cartilage and bone because of the more incoherent orientation maps.

**Figure 3:**
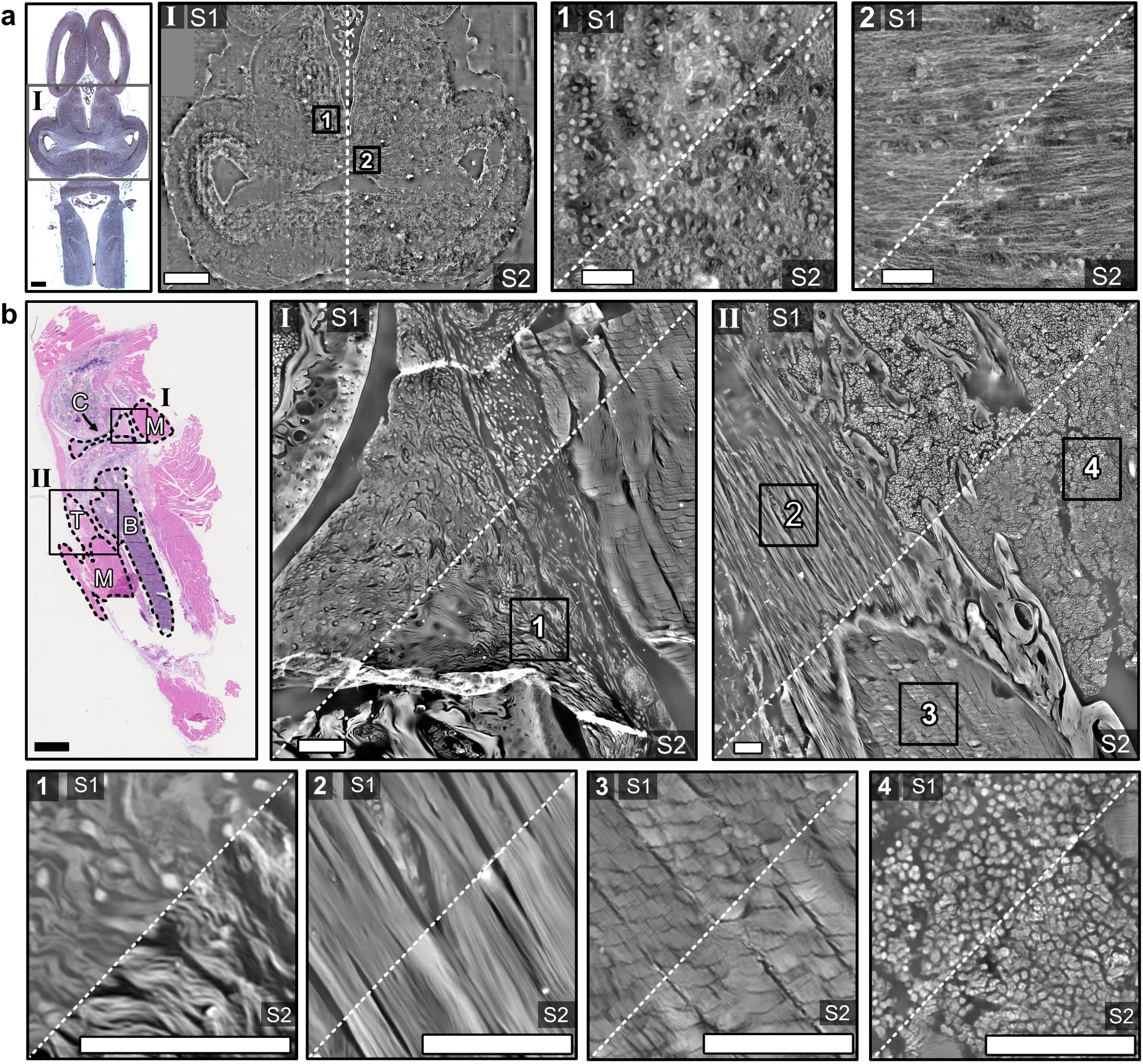
Comparison of the phase maps obtained from both setups (S1 and S2), in radians. a) Left: Zoomed-in middle region of the frog brain section (I); scale bars indicate 1 mm. The top left corner was not scanned with setup S1 and filled with zeroes to match the field of view of setup S2. Right: Zoomed-in region showing cell nuclei (1), zoomed-in region of horizontal nerve bundles (2); scale bars correspond to 100 µm. b) Top: Zoomed-in regions of the mouse knee section containing different tissue types (M–muscle, C–cartilage, T–tendon, B–bone). Bottom: Zoom-ins of the four marked regions at the top showing details of cartilage (1), tendon (2), muscle (3), and bone (4) tissue. Scale bars correspond to 100 µm.

### 2.2 Reconstructed high-resolution phase images

**Figure 3** shows the FPM phase maps obtained from both setups, S1 and S2, for the frog brain section (a) and the mouse knee section (b). The zoomed-in regions feature different tissue types: neuronal cell bodies (a1), nerve fibers (a2), cartilage (b1), tendon (b2), muscle (b3), and bone (b4). The contrast of the phase maps is proportional to the optical thickness of the sample which is related to its refractive index and density, allowing to quantitatively image the sample’s morphology and provide endogenous contrast to the specimen images.

The reconstructed structures in the displayed phase maps are consistent to each other for both setups and show similar details with slightly different contrast. The resolution is slightly higher for setup S1, as expected. Residual low-frequency artifacts in S2 stemming from the non-telecentric configuration were compensated as described in the Methods section. It is remarkable that setup S2 can produce these high-quality phase maps although the setup was not intentionally designed for FPM.

### 2.3 Combined high-resolution phase and fiber orientation maps

**Figure 4** shows the FPM phase maps overlaid with the ComSLI fiber orientation maps for some selected tissue regions for setup S1 (a-c) and setup S2 (d). The multi-modal images convincingly show the additional information that is obtained by Fourier Ptychographic Scattered Light Microscopy (FP-SLM). The higher-resolution phase maps allow for more structures to be shown alongside the orientation maps. The color-overlaid images were generated by upscaling the orientation maps to the high-resolution reconstructions, keeping the saturation and value, but setting the hue according to the direction map color. For some zoomed-in regions, the high-resolution phase maps were overlaid with vector distribution maps, which show the orientations as colored lines for better visibility. In these combined high-resolution phase and fiber orientation/vector maps, the direction coherence of any fiber bundle becomes immediately evident. The zoomed-in thigh muscle region (bI) nicely shows the stripe-structure in skeletal muscle (alternating between red/magenta and green/blue stripes). This structure is invisible when looking at the FPM phase map alone (bottom triangle in b1). The complex structure of tissues such as cartilage becomes directly apparent when looking at the combined map (cI and c1). As there is no additional information necessary to perform ComSLI analysis on a dataset that has originally been used for FPM analysis, these color and vector overlays can be generated alongside the high-resolution (HR) reconstructions, and can serve as an additional information overlay onto the results that adds fiber bundle specificity to FPM. On the other hand, high-resolution phase maps add a level of morphological detail to ComSLI fiber orientation maps, which naturally have a lower resolution and only indirectly image-oriented structures. The phase maps put the computed orientations into histological context, as can be appreciated when looking at zoom-ins of tendon, bone, and muscle tissues (d). The vector distribution map (d2) shows the crossing substructure of muscle tissue that becomes visible with setup S2, as already discussed for Figure 2g. The results of Figure 4 further show the complementarity between FPM and ComSLI which reinforce each other in a consistent way, thus showing the strengths of FP-SLM.

### 2.4 Quantitative comparison to structure tensor analysis

To quantitatively assess the performance of FP-SLM, the ComSLI fiber orientations obtained from both setups were compared with those obtained by applying STA to the FPM phase maps (see Methods). **Figure 5** shows the pixel-wise comparison of the fiber orientation maps as split-views (b,e) and as difference maps with corresponding histograms (c,f) for a region in the frog brain (nerve fiber bundle) and a region in the mouse knee (M–muscle, T–tendon, B–bone), indicated by the rectangles in the bright-field microscopy images (a,d). As can be seen from Figure 5, the ComSLI fiber orientations are overall in good correspondence to the structures obtained from the STA; the histograms show distributions with mode and mean near zero for the selected regions of interest (see Supplementary Table 2 for the complete metrics). In regions with less aligned structures such as brain gray matter (c, top) and bone (f, top right), larger absolute differences are observed as expected.

**Figure 4:**
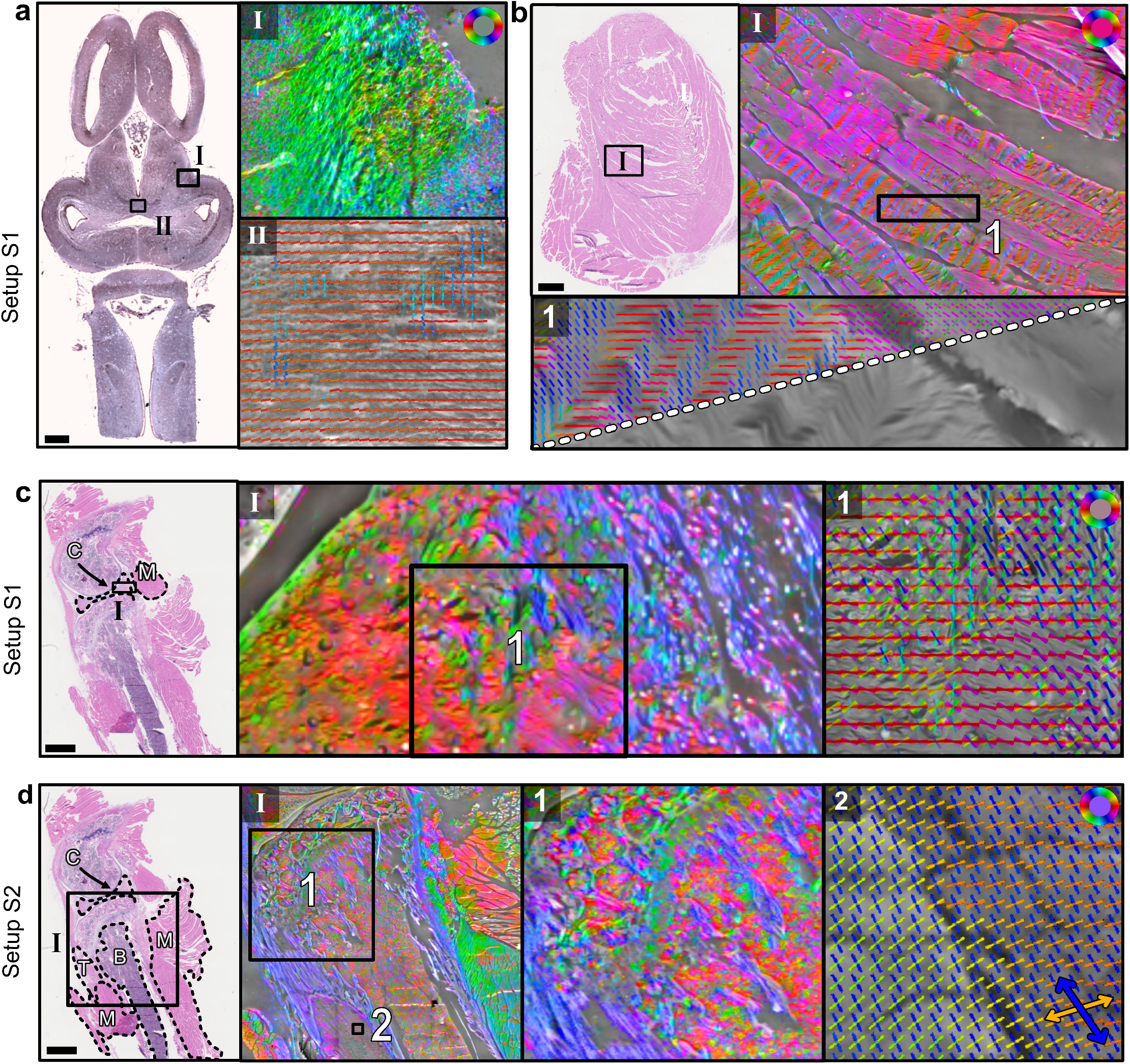
FPM phase maps overlaid with ComSLI fiber orientation and vector distribution maps for setup S1 (a-c) and S2 (d), shown for different selected regions, as indicated by the rectangles in the corresponding bright-field microscopy images on the left of the respective panels. The fiber orientation maps show the color-coded orientations for each ComSLI image pixel according to the color wheel in the top right of each panel. Vector distribution maps show orientations as colored lines for kernels of x × x pixels overlaid. a) Zoomed-in regions of the frog brain section showing nerve fiber bundles. The top right image (I) shows the fiber orientation map, the bottom right image (II) the vector distribution map (x = 15) overlaid with the respective FPM phase map. b) Zoomed-in regions of the mouse thigh muscle sample. The top right image (I) shows the phase map overlaid with the fiber orientation map, the bottom image (1) shows a more detailed zoom-in of the phase map with/without overlaid vector distributions (x = 10) in the top/bottom triangle for an area with visible muscle-disk stripes. c) Zoomed-in region of the mouse knee sample containing cartilage (C). The left image (I) shows the fiber orientation map, the right image (1) the vector distribution map of a zoomed-in region (x = 20) overlaid with the corresponding phase map. d) Zoomed-in regions of the mouse knee sample containing cartilage (C), muscle (M), tendon (T), and bone (B). Region (I) shows the phase map overlaid with the corresponding fiber orientation map. The more detailed zoom-in (1) shows the fiber orientations in tendon and bone tissue, while (2) shows the vector distributions (x = 10) in muscle tissue featuring the crossing substructure that becomes visible with setup S2. Scale bars correspond to 1 mm.

**Figure 5:**
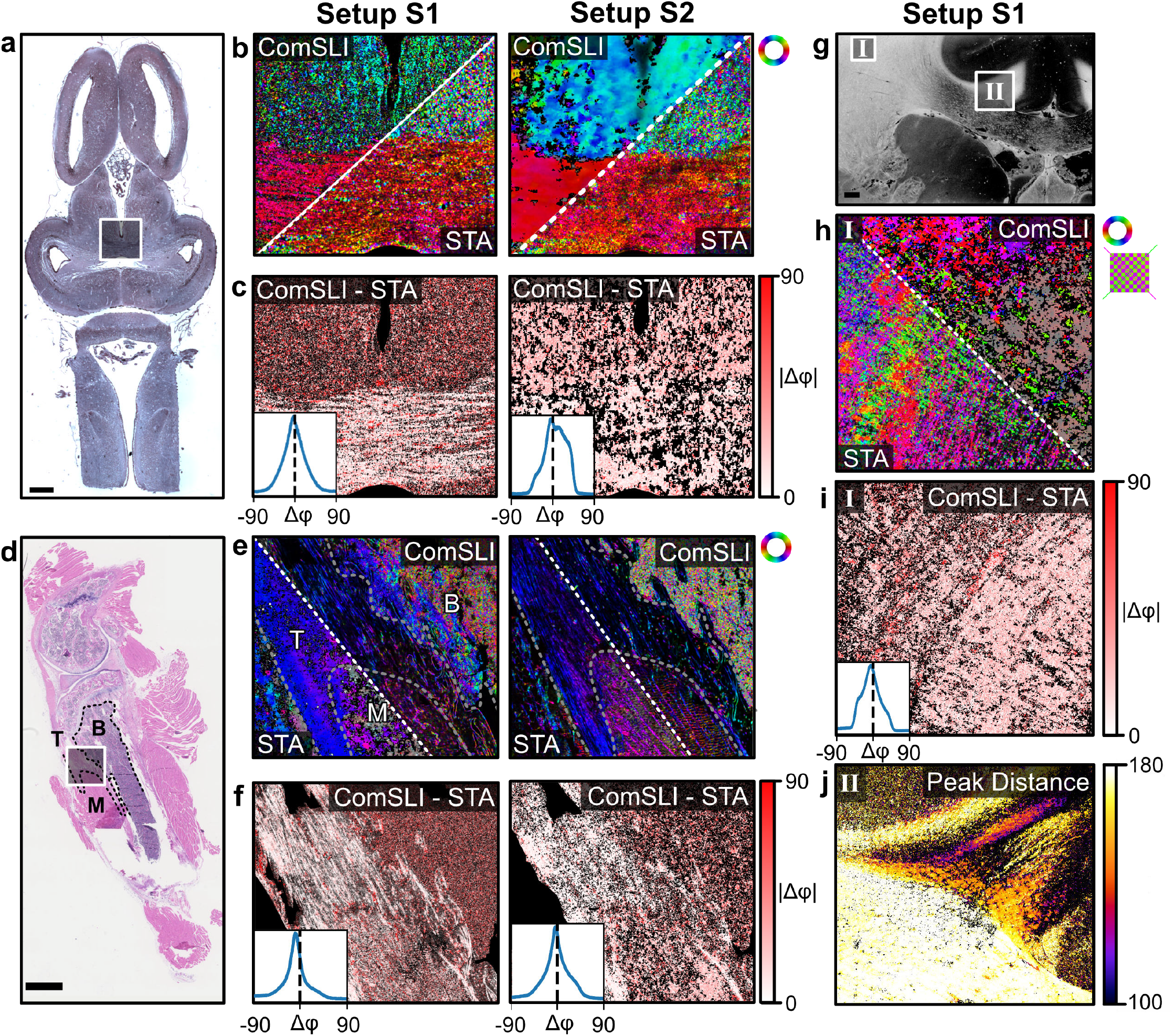
Quantitative comparison of ComSLI fiber orientation maps and those obtained from STA analysis on the corresponding FPM phase maps for both setups. a),d) Bright-field microscopy images of the frog brain (top) and mouse knee section (bottom). M–muscle, T–tendon, B–bone. b),e) ComSLI fiber orientation maps (top triangles) obtained from measurements with setup S1 (left) and setup S2 (right), shown together with the corresponding STA fiber orientation maps (bottom triangles) for the regions indicated by the white rectangles in (a) and (d). c),f) Difference between the ComSLI and STA fiber orientation maps shown as difference map (absolute difference) together with the corresponding histograms (inserts) for the selected regions, in degrees. g) Average scattering signal of a vervet monkey brain section obtained from a measurement with setup S2 [26]. h) ComSLI (top triangle) and STA (bottom triangle) fiber orientation maps of a crossing fiber region (corona radiata, area I in g). ComSLI yields dual-colored pixels indicating two crossing fiber orientations (see zoomed-in region with green and magenta arrows), while STA yields a single fiber orientation per pixel. i) Corresponding difference map (absolute difference) and histogram (insert). j) ComSLI peak distance map of a steep fiber region (cingulum, area II in g), in degrees. Histogram metrics (mean, mode, full width at half maximum, root mean square error) can be found in Supplementary Table 2. Scale bars correspond to 1 mm.

To study the effect of crossing fibers, a coronal vervet brain section (cf. Supplementary Figure 1c, left) has additionally been imaged with setup S2 as shown in previous publications [26], and compared in the same way for a region in the corona radiata (Figure 5g, region I) which contains many crossing nerve fibers. While the ComSLI fiber orientation map (h, top triangle) shows the multiple crossing fibers per image pixel (dual-colored pixels as shown in the zoom-in on the right), the STA fiber orientation map (h, bottom triangle) only reveals a single orientation per pixel, leading to larger differences in the difference map and histogram (i).

Another difference is that the ComSLI analysis can be used to gain information about the out-of-plane fiber orientation. Previous studies have shown that the high-intensity line in the scattering pattern of a fiber bundle becomes curved with increasing out-of-plane angle of the fiber bundle. In the azimuthal line profile, in which the intensity values of a scattering pattern are plotted against the azimuthal angle (cf. Supplementary Figure 4), this curvature can be measured as the distance between two peaks. Figure 5j shows the peak distance for each image pixel in region II of the vervet brain section: regions with in-plane fibers have a peak distance of approximately 180 degrees, smaller peak distances (in the upper right corner) correspond to more inclined fibers; this region is the cingulum in the vervet brain, as shown in previous publications [22, 26]. While ComSLI provides 3D structural information, STA analysis only yields 2D information; FPM (in the absence of tomographic inversion algorithms) provides an integral information content.

## 3 DISCUSSION

We introduced Fourier Ptychographic Scattered Light Microscopy (FP-SLM). Taking advantage from the fact that Fourier Ptychographic Microscopy (FPM) and Computational Scattered Light Imaging (ComSLI) use the same basic setup (Figure 1a), we combined for the first time FPM and ComSLI analysis on the same dataset. We show that ComSLI analysis can be applied to an image stack originally used for FPM in order to reveal intricate fiber organization (Figure 2), adding relevant structural information and the possibility to trace fiber pathways in dense tissues. Vice versa, we show that FPM processing can be applied to an image stack originally used for ComSLI in order to reconstruct high-resolution images of the measured tissue sample (Figure 3), with 5 × higher resolution than maps obtained from ComSLI analysis alone. The results are comparable to ComSLI fiber orientation and FPM phase maps obtained from state-of-the-art setups that were originally used for FPM (setup S1) and ComSLI (setup S2) alone, showing that FPM and ComSLI analyses are compatible with each other. Setup S1 has originally been designed for high-resolution, sub-cellular analysis, and therefore has a more than five times higher magnification than setup S2. Setup S2 has originally been designed for disentangling interwoven fiber pathways over large fields of view, and thus employs a larger LED array than setup S1 (176 × 176 vs. 32 × 32 LEDs, see Methods) to allow for a more detailed reconstruction of scattering patterns. For the generation of ComSLI fiber orientation maps, we employed a new filtering approach (see Methods) which allows for a more detailed analysis of fiber orientations and provides a visible improvement over the previous approach without filtering (see Supplementary Figure 4, a5 vs. b5), which will also improve future ComSLI analyses.

Looking at the phase maps obtained from both setups (Figure 3), there are some differences in resolution and synthetic NA: Setup S2 employs weaker LEDs than setup S1, and they are located further away from the sample, which puts a limit on the dynamic range. In order to keep the measurements with central LEDs from overexposure, there is an outer limit of the LED polar angle of around 35° with which measurements can be made (see Methods). This places a limit on the final synthetic NA. Other studies have overcome this by setting different exposure times for different measurements, then adjusting intensity values during post-processing [27].

FP-SLM enables the generation of combined high-resolution phase and fiber orientation maps (illustrated for various tissue types in Figure 4), unlocking new possibilities for analyzing biological samples. Thanks to this novel approach, it is possible to reveal subtle morphological tissue traits and the fiber structural organization in a single image with multiple information channels.

As both FPM and ComSLI are label-free techniques, they can be used to examine histological samples in their native state, without the need for fluorescent dyes or stains that may alter the properties of the tissue. This is particularly beneficial for retrospective studies, where archived tissue samples can be reanalyzed.

In this study, we compared the fiber orientation maps obtained from ComSLI analysis to results obtained from structure tensor analysis (STA) applied to the corresponding FPM phase maps (Figure 5). Besides the mere qualitative inspection of the images, a pixel-wise quantitative comparison shows that the reconstructed fiber orientations are similar to each other, which validates our approach. At the same time, STA applied to phase maps obtained from FPM cannot replace the ComSLI analysis. STA cannot distinguish between low- and high-scattering structures, and needs sufficient structural detail in the realspace image to reconstruct directed structures, and performs better in sparsely populated tissues than in regions with densely packed fibers like brain white matter. In crossing fiber regions like the corona radiata (cf. Figure 5h), standard STA falls short in resolving multi-directional fibers. ComSLI on the other hand provides multiple fiber orientations per image pixel, and it allows reconstructing scattering patterns from the raw data (cf. Supplementary Figure 3f). This has the advantage that additional information like 3D fiber orientation, fiber size, or tissue type could be extracted. While extensions to standard FPM allow for 3D tomographic phase microscopy [28], the setup and measurement are significantly more complex. The standard FPM and STA analyses presented here only yield 2D information and no specificity for different fiber types. ComSLI has the potential to estimate out-of-plane fiber orientations from the scattering peak distance [22, 26] as shown in Figure 5j, thus adding 3D orientational information to the 2D results from the FPM and STA analyses applied here. At the same time, applying STA to the FPM phase maps yields higher resolution and can complement the ComSLI fiber orientation map. On a general note, STA applied to FPM phase maps yields better results than STA on standard brightfield microscopy images. Schurr et al. applied STA on Nissl-stained brain sections in order to reconstruct their nerve fiber organization [29]. In that work, specific cell staining was required to indirectly reconstruct the course of the nerve fibers as the contrast was not sufficient for STA. Using either FPM and subsequent STA, or ComSLI, it becomes possible to reconstruct fiber orientations also in unstained tissues that exhibit little contrast.

Avoiding the use of stains is advantageous for several reasons. In the first place, the measurement is non-invasive and does not alter the quantities to be measured. Cytotoxicity effects linkable to the use of dyes are widely reported. Photobleaching is another issue to be considered since it can generate ambiguities in the interpretation of phenomena occurring over long time scales, e.g., during time lapse experiments. Above all, the use of stains to enhance the contrast of tissue slides may generate ambiguous results since the appearance of the images may depend on the staining protocol applied, the lab environment, as well as the lab operator [30]. The application of ComSLI analysis to data that was originally acquired for FPM can be an interesting future pathway to access specific information from non-specific quantitative phase FPM acquisitions of stain-free tissue slides.

Our work is a first proof-of-concept study, showing the compatibility of both FPM and ComSLI analyses and the broad applicability across different species (mouse, frog, vervet) and types of tissue (muscle, cartilage, bone, tendon, brain). The compatibility of the two different techniques and their cross-analysis are not obvious and have never been demonstrated so far. Our developed analysis software for FP-SLM can easily be adapted to setups from other groups that were originally designed for FPM or ComSLI (consisting of an LED array and low-NA objective) and be used to compute additional fiber orientation or high-resolution phase maps from their existing datasets.

We have demonstrated FP-SLM on state-of-the-art FPM and ComSLI setups (setup S1 and S2), and quantitatively compared their performances. Notably, each investigated system can stand on its own, i.e., both FPM and ComSLI can be performed using setup S1 or S2, without need to design and build an extra FP-SLM setup. As our study has shown that FP-SLM works for both extremes, i.e., a setup optimized for FPM and a setup optimized for ComSLI, any system trying to establish a compromise, e.g., between the FPM resolution and ComSLI FoV, will have a performance spanning in the middle between these two state-of-the-art configurations. Within the large plethora of possible imaging choices for FP-SLM setups, we have presented and analyzed two different examples of integrated systems that represent two extremes of design trade-off choices, which depend on the specific imaging needs. Any other design solution between these two extremes is in principle viable to build a new FP-SLM apparatus. The reader can choose the preferred configuration based on the respective research question and individual requirements, i.e., whether a large field of view or a high optical resolution is preferred, and how much time should be spent on measurements and image reconstruction. While setup S1 yields high-resolution phase maps and reliable fiber orientation maps for small fields of view, setup S2 provides a larger field of view in the same measurement time and more detailed scattering patterns, but at the cost of a slightly lower resolution in the reconstructed phase maps.

Our work opens up new lines of research and has an especially big impact for retrospective studies. So far, ComSLI has only been used to analyze fiber organization. With FP-SLM, we can apply the adapted FPM analysis to any existing ComSLI dataset to generate a high-resolution real-space image and a higher-resolution fiber orientation map (through subsequent STA). The quantitative tissue morphology provided by the phase map removes the limitation of ComSLI to indirectly image fibers and helps to put the fiber orientation map better into context, greatly simplifying the analysis of histological tissues. ComSLI has been shown to provide similar fiber orientation maps as polarization microscopy, with the advantage that it yields multiple fiber orientations per measured image pixel [22] and that it also works on non-birefringent samples independent of sample preparation [19]. While ComSLI is a relatively young technique, polarization microscopy has been developed more than a century ago to make directed tissue structures visible, and is an established imaging method in research and clinical laboratories for analyzing histological tissue samples. Up to now, research groups had to make a choice whether to focus on high-resolution or directional information when designing an imaging system. Our work finally enables the Fourier Ptychography community to use their FPM datasets to generate fiber orientation maps that are similar to those obtained with polarization microscopy – without the need to perform additional measurements or to purchase an additional setup, and providing fiber orientations also in regions with multiple crossing fibers and in non-birefringent specimens. With FP-SLM, any existing FPM dataset can now be re-processed with an adapted ComSLI analysis to generate a fiber orientation map and analyze the structural tissue organization of the measured sample. No changes in the existing imaging apparatus are needed, and the additional parameter maps are by design perfectly registered to the original maps, enabling pixel-wise analysis. As FP-SLM can be applied to any existing FPM or ComSLI dataset, it will greatly extend the possible applications of both Fourier Ptychography and Computational Scattered Light Imaging, and enable retrospective studies of archived tissue samples, e.g., analyzing the fiber organization in biological samples.

## 4 MATERIALS AND METHODS

### 4.1 Sample preparation

The mouse samples were obtained from a male nude mouse (strain: Rj:NMRI-Foxn1nu/nu, Janvier Labs). The mouse was sacrificed at 5 months using cervical dislocation. The leg was removed, fixed in 4% formaldehyde, dehydrated in increasing alcohol series (70%, 80%, 90%, 96%, 100% ethanol), treated with xylene, embedded in paraffin, and cut longitudinally with a microtome (Leica RM2165) into 4 µm-thin sections. The sections were placed in a decreasing alcohol series to remove the paraffin, mounted on glass slides, stained with hematoxylin and eosin (H&E), and cover-slipped. Two sections from the middle, one containing only thigh muscle, and one containing also bone and knee joint (see Supplementary Figure 1b), were selected for further analysis and measured about 4 months after preparation. All animal experiments were approved by the Animal experiment committee under the Dutch experiments on Animal Act, with license reference number AVD10100-2216075, and following ARRIVE guidelines.

The frog brain is part of the Archive Collection of the Istituto di Cibernetica, Neuroanatomy Laboratory, National Research Council, Italy [31]. It was obtained from a male tadpole and prepared in a similar way as the mouse samples; after paraffin embedding, the brain was horizontally cut into 10 µm-thin sections, deparaffinized, and Golgi-stained; a section from the middle (see Supplementary Figure 1a) was selected for further analysis.

The vervet monkey brain was obtained from a healthy 2.4-year-old adult male in accordance with the Wake Forest Institutional Animal Care and Use Committee (IACUC #A11-219). Euthanasia procedures conformed to the AVMA Guidelines for the Euthanasia of Animals. All animal procedures were in accordance with the National Institutes of Health guidelines for the use and care of laboratory animals and in compliance with the ARRIVE guidelines. The brain was perfusion-fixed with 4% paraformaldehyde, removed from the skull within 24 hours after death, immersed in 4% paraformaldehyde for several weeks, cryo-protected in 20% glycerin and 2% dimethyl sulfoxide, deeply frozen, and coronally cut from the front to the back into 60 µm-thick sections using a cryostat microtome (Polycut CM 3500, Leica Microsystems, Germany). The brain sections were mounted on glass slides, embedded in 20% glycerin, and cover-slipped without any staining. A section from the middle (no. 505) was selected for further analysis (see Supplementary Figure 1c). The section was embedded in 2012, i.e., 12 years before the measurements of this study.

### 4.2 Bright-field microscopy

The brain section of the frog tadpole (as shown in Figure 2a and other figures) was measured with setup S2 with all LEDs turned on, with 2.87 µm pixel size and 0.83 × magnification. The tissue sections of the mouse leg (as shown in Figure 2e,h and other figures) were measured with Hamamatsu Photonics K.K.’s Nanozoomer 2.0 HT digital slide scanner to scan the H&E-stained microscopy slides at 20× magnification with 0.46 µm pixels.

### 4.3 Setup S1

The setup has originally been designed for FPM analysis [24, 25]. It is sketched in Supplementary Figure 2a. It consists of a planar 32 × 32 RGB-LED matrix (Adafruit Industries), with 4-mm distance between two adjacent LEDs. The distance between the central LED and the sample plane is 4.67 cm. In total, 177 images were collected, corresponding to a circle with 15-LED diameter (cf. Supplementary Figure 3a), i.e., the farthest LED from the center of the matrix is positioned at 28 mm distance and illuminates the sample at an angle *θ*_max_ ≅ = 31°. Red LED emission at 632 nm central wavelength was used for illumination. The setup contains a 4 × achromatic Plan N microscope objective (Olympus) with numerical aperture NA_obj_ = 0.1, and a 400 mm tube lens that redirects light towards the camera (CCD, Photometrics Evolve 512, 4.54 µm pixel pitch). The overall optical system provides a 4.29 × magnification on the camera plane that coincides with the sample best focus plane [25]. Each intensity image was captured with an exposure time of 800 ms, quantized with 12-bit depth, and consists of 1940 × 1460 square pixels with a pixel size in the image plane of 1.06 µm, corresponding to a field of view of 2.05 × 1.55 mm^2^; the lateral optical resolution is at least 2.76 µm for the low-resolution images. The resulting lateral optical resolution after applying the synthesis of all the images is at least 0.49 µm (see below), in good agreement with the theoretical prediction. The lateral optical resolution has been verified by acquiring the image of an USAF target in through-transmission as reported in [32].

### 4.4 Setup S2

The setup has originally been designed for ComSLI analysis [23]. It is sketched in Supplementary Figure 2d. It consists of an LED panel (Absen LED AW2.8), placed 15 cm underneath the sample holder. This panel is square with 176 × 176 RGB-LEDs arranged in a grid pattern, with 2.87 mm of LED pitch. It features red (630 nm), green (520 nm) and blue (470 nm) LEDs, all with a bandwidth of 20 nm. For the measurements shown here, the mouse knee sample was imaged with green light, and the frog brain, vervet brain, and mouse muscle samples with blue light. The samples were placed underneath an optical system containing a lens (Rodenstock Apo-Rodagon-D120, numerical aperture: NA_obj_ = 0.045, focal length: 120 mm), and an RGB camera (Basler acA5472-17uc) with a Bayer filter. The lens was adjusted to an f-number of 5.6, and positioned 264 mm away from the sample, yielding a magnification of 0.83. A large styrofoam funnel was attached to the lens to block out any errant light that is not coming from the sample. Additionally, any reflective part of the setup was taped over with black diffusive tape. During a measurement, multiple acquisitions were made: during each acquisition, a single LED was lit on the panel, the sequence of which follows a raster-scan pattern, across a square of 40 × 40 LEDs, such that the maximum polar angle that is measured in all directions is 20°. Each time an LED lights up on the LED panel, 8 images with 300 ms exposure time were acquired, and summed together to create one image. The acquired images contain 5472 × 3648 12-bit square pixels with a pixel size of 2.87 µm, corresponding to a field of view of 1.57 × 1.05 cm^2^; the lateral optical resolution determined with an USAF target using blue light is at least 4.38 µm. These images were de-interlaced and saved as 16-bit integer RGB tiff files.

### 4.5 High-resolution phase reconstruction

The idea behind FPM is to enhance the resolution beyond the limit of the optical system by adding a synthetic illumination numerical aperture (NA_ill_) to the low-NA microscope objective (NA_obj_) [5, 33]. The sample is illuminated from *N* different angles by sequentially switching on LEDs in an array. A set of bright- and dark-field intensity images is collected and processed together to provide the HR complex amplitude of the sample by applying iterative phase-retrieval algorithms [5, 34, 35]. In the low-NA configuration, this approach gathers amplitude and phase-contrast images with large space-bandwidth product (SBP), i.e., optimizing at the same time field of view and lateral resolution. Each of the acquired images collects a different portion of spatial frequencies of the sample. In particular, the images collected in dark-field illumination mode are responsible for synthesizing the HR details that would be lost otherwise since the corresponding signal falls outside the limited NA_obj_. The iterative reconstruction process is based on a joint optimization for the object complex amplitude, *o*, and the system impulsive response, *h*, from the set of *N* intensity measures. Let 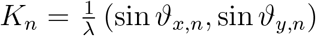, *n* = 1, …, *N* be the *n*-th illumination vector, modeled as a plane wave impinging from the direction defined by the couple (*ϑ*_*x,n*_, *ϑ*_*y,n*_). The corresponding low-resolution (LR) intensity image captured by the camera takes the form:

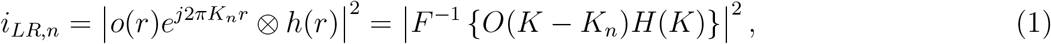

where *o* is the object complex amplitude, *r* represents the spatial coordinates vector, ⊗ and *F* {…} are the convolution and 2D Fourier transform operators, respectively, and we use the capital letters to refer to the Fourier domain. *H* is a circular pupil with radius determined by NA_obj_*/λ*. In other words, each measurement is a sample in the image plane of the object spatial frequencies *O*(*K* − *K*_*n*_) filtered through the system’s optical transfer function *H*(*K*) and the sensor that operates the square modulus (thus losing the phase information). The phase retrieval process performs an inner loop using a gradient descent method to iteratively optimize both the complex amplitude and the phase in the frequency domain. The whole process is then repeated *M* times until guaranteeing a stable solution for both the HR complex amplitude and the system impulse response. Denoting with *K* = [*K*_1_, …, *K*_*N*_] the set of illumination vectors and with Γ{…} the functional that describes the whole iterative FPM process summarized above, the HR phase-contrast map is obtained as 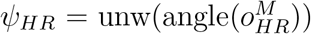, where unw(…) is the phase unwrapping operator [36] and 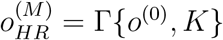, is the estimated complex amplitude after *M* iterations.

Phase-contrast maps show the contrast in the optical delay experienced by light transmitted through the specimen with respect to its unperturbed propagation through the surrounding medium. This delay, expressed in radians, is proportional to the optical thickness of the specimen, i.e., the product between its refractive index and its physical thickness. In most FPM systems, the phase-contrast map is the final outcome, allowing direct imaging of the specimen’s morphology. In this sense, the phase-contrast map is quantitative, since each pixel is directly associated to a physical sample parameter. In order to decouple the refractive index and the physical thickness, tomographic approaches [28] are needed which estimate the 3D refractive index distribution of the specimen. The dry mass is proportional to the refractive index and is a measure of the density of the specimen in each specific pixel (e.g., the nucleus of a cell has a higher dry mass with respect to the cytoplasm). While bright-field images show poor contrast for stain-free samples, phase-contrast maps show high contrast which is why FPM and other quantitative-phase-imaging methods work on stain-free slides.

### 4.6 FPM phase maps (setup S1)

Setup S1 employs a microscope objective with NA_obj_ = 0.1 and a maximum illumination angle of 31°. This configuration creates a synthetic numerical aperture of: NA_synth_ = NA _obj_ + NA_ill_ = 0.1 + sin(31°) ≈ 0.6. The iterative scheme that was used to generate the HR phase maps from the images measured with setup S1 is similar to the well-known Gerchberg-Saxton algorithm, where at each iterative step the guess of the HR object is updated using the measured intensity as a constraint while leaving the phase unaltered. The system pupil is updated accordingly. At the *n*-th iteration, the portion of the HR spectrum centered in the coordinates set by the spatial frequency vector *K*_*n*_ is replaced using the *n*-th intensity measure acting as a constraint. The algorithm iterates between the spatial and the frequency domains by replacing the measured LR amplitude only, while the phase keeps memory of the process and is allowed to converge by minimizing a cost function [35, 37]. Each image recorded with setup S1 was divided into overlapping patches with size 100 × 100 pixels and 30 pixels overlap. The EPRY [5, 35, 37] iterative algorithm described above was applied separately to each patch. The main driving reason behind this choice was to allow faster processing through a highly parallelizable process and, above all, to make the FPM assumption of plane wave illumination valid [5]. After applying the FPM reconstruction of the high-resolution complex amplitude, each patch sizes 500 × 500 pixels. The HR amplitude and phase-contrast images were stitched together using an alpha blending algorithm. The resulting reconstruction is a 9500 × 7000 pixel image with 0.21 µm pixel size, at least 0.49 µm lateral optical resolution, and 2.05 × 1.55 mm^2^ field of view, i.e., with a space-bandwidth product of 6 · 10^6^.

### 4.7 FPM phase maps (setup S2)

Setup S2 employs an objective with NA_obj_ = 0.045 and a maximum illumination angle of 20◦. This configuration creates a synthetic numerical aperture of: NA_synth_ = NA_obj_ + NA_ill_ = 0.045 + sin(20°) ≈ 0.39. To create phase maps from the collected images, the open source FPM library called PtyLab [38] was used. This library takes a stack of measurement images as input, along with the measurement metadata and a number of hyperparameters. The PtyLab qNewton engine was used for each reconstruction, going through 100 iterations. Each image was divided into overlapping patches, with a patch size of 128 × 128 pixels and an overlap of 32 pixels on each side. The patches were up-scaled before reconstruction to 1024 × 1024 pixels using bicubic interpolation. The size of the patches determines the time required to reconstruct the final phase map. Due to the non-telecentric optics of setup S2, there are still significant phase-curvature artifacts expected in the final reconstruction. These were anticipated by multiplying the upscaled patches with the expected phase curvature before reconstruction, and repeating this multiplication after the reconstruction [39]. We here decided to use a relatively small patch size compared to the large field of view of setup S2 in order to ensure that the plane-wave assumption is largely valid and the correction of the phase curvature only has to be minor. After completed reconstruction, the phase map was obtained by taking the angle of the transformed result, and unwrapping the phase using a least squares method [36] to correct for phase wrapping in optically thick tissue regions. Subsequently, a Gaussian low-pass filter (*σ* = 50 pixels) was applied to each patch in order to estimate a reference background map for each patch. Similarly to the process of phase aberration compensation using a reference hologram that is typical for digital holography, the reference background maps were subtracted from the unwrapped phase maps in order to remove residual low-frequency artifacts caused by the non-telecentric optics. This process also promotes a more effective stitching of the patches, which was realized using alpha blending [40] to obtain the final phase map. The maps have a pixel size of 0.56 µm; the lateral optical resolution determined with an USAF target using blue light is at least 0.87 µm, corresponding to a 5× increase in resolution compared to the images obtained from a ComSLI analysis alone.

### 4.8 Scattering pattern generation

In ComSLI analysis, fiber orientations are computed from scattering patterns (cf. Figure 1, d1). To compute these scattering patterns, the measured images from both setups were first saved to a 4D array *I*(*x, y, u, v*) where (*x, y*) refers to the pixel index in the field of view, and (*u, v*) refers to the index of the LED array. A scattering pattern can be obtained by selecting one (*x, y*) coordinate, and plotting the intensity values according to the (*u, v*) coordinate. Each scattering pattern then represents the intensity response at a single pixel while changing the illumination angle. Spatial position and illumination angle are not entirely uncoupled though. The illumination angle varies slightly across the field of view. This also has an effect on the scattering patterns, namely that the (*u, v*) coordinates which describe direct illumination by an LED shift linearly depending on the (*x, y*) coordinates. Assuming that the (*x, y*) coordinates are centered around the optical axis, and that the (*u, v*) coordinates have their origin in the center of the LED board, the scattering pattern from the pixel in the middle of the field of view should have its direct illumination in the middle. The subpixel shift of this point can be calculated with:

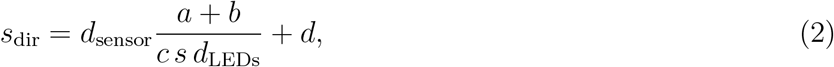

using the camera pixel pitch *d*_sensor_, the distance between LEDs *d*_LEDs_, the kernel stride *s*, and the distances between display and sample *a*, sample and lens *b*, and lens and camera *c*. An additional correction term *d* can be calibrated, which describes the distance of the LED panel center to the optical axis of the camera. Once the (*u, v*) coordinates have been adjusted according to the direct illumination, the illumination angle can be calculated on every pixel in the scattering pattern. These angles, combined with the setup NA, provide the coverage in frequency space from which these scattering patterns sample their data. Supplementary Figure 3a-d shows the LED arrays and the coverage in frequency space both for setup S1 (left) and setup S2 (right). The bottom part of the figure shows the reconstructed scattering patterns for three selected locations in the mouse knee sample, showing the good correspondence between the scattering patterns obtained from both setups.

### 4.9 ComSLI fiber orientation maps (setup S1 & S2)

ComSLI analysis rests on the principle that light encountering an elongated object tends to scatter more in the direction perpendicular to the object orientation [41]. Hence, by determining the direction of high scattering, the direction of aligned structures like nerve, muscle or collagen fibers can be determined. For this purpose, an azimuthal line profile is generated from the scattering patterns to better determine the positions of high-scattering peaks and thus the fiber orientations. While previous works sample the scattering pattern along a certain azimuthal angle, we used here for the first time a filter approach to measure the intensity along each azimuthal angle, see Supplementary Figure 4. The filter was separated into an azimuthal part and a radial part, as inspired by the work of Kittisopikul et al. [42]. They describe a combination of an annular and orientation filter in the frequency domain in order to extract accurate orientations from filamentous images, even in the presence of junctions. The scattering patterns were directly treated as the frequency domain of the local area around their pixel. Before filtering, a logarithm was applied to the scattering patterns. Pixels in the center of the pattern have intensities that are orders of magnitude higher than pixels in the outer regions, yet it is this signal in the outer regions that is most important. An annular filter (Supplementary Figure 4, b1) was used to bring even more attention to these outer regions, described as:

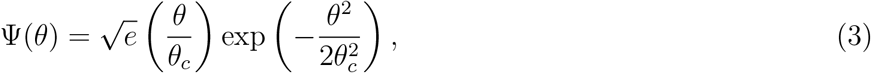

where *e* is Euler’s number, *θ* is the polar spatial component (in radians), and *θ*_*c*_ the central polar angle of the filter. The azimuthal part of the filter was implemented with a von Mises distribution (Supplementary Figure 4, b2), described as:

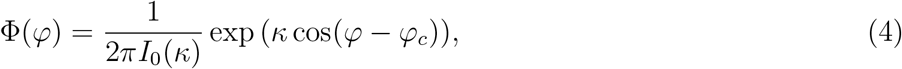

where *ϕ* is the azimuthal spatial component, *ϕ*_*c*_ the central azimuthal orientation, *κ* a constant which determines the spread of the filter, and *I*_0_ is the Bessel function of the first kind of order 0. The central polar angle of the annular filter was set to *θ*_*c*_ = 1.57 rad^−1^, and *κ* was set to 30, these values were empirically derived. For each pixel in the data stack, its spatial (*x, y*), and LED index (*u, v*) coordinates were used to calculate the spherical coordinate angle (*θ, ϕ*) of that pixel to the LED, resulting in an array *I*(*x, y, θ, ϕ*), where each spatial coordinate (*x, y*) has an intensity profile described by the angular co-ordinates (*θ, ϕ*). The above described filters were then used to calculate the spatial coordinate’s intensity response along an azimuthal direction:

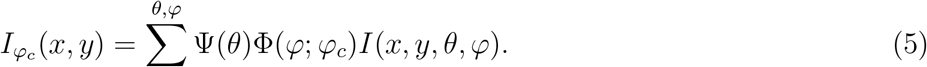

This process was repeated for 24 equally spaced azimuthal orientations *ϕ*_*c*_, spanning a full circle, resulting in a filtered image stack *I*(*x, y, ϕ*_*c*_). Because *θ* and *ϕ* were not sampled uniformly (cf. Supplementary Figure 3c-d), an additional calibration image stack was created using the same coordinates as the image stack, but all intensity values are constant. Each value in the filtered image stack was divided by its corresponding value in the calibration stack. Each pixel in (*x, y*)-space in the 24-layer image stack contains in the *ϕ*_*c*_-direction a line profile, representing the sample response per angle (Supplementary Figure 4, b4). This line profile contains two peaks per prominent direction in the sample structure at that pixel, spaced 180° ± 35° apart for in-plane fibers. The Scattered Light Imaging ToolboX (SLIX) [22, 43] was used to identify these peaks, refine their position by calculating the geometric center of the peak tip, and assign peak pairs based on angular distance. The orientation of the structure was then determined by the mid position of a peak pair (red dashed line in b4). SLIX can identify up to three fiber orientations this way, and represents them in a fiber orientation map (FOM), where each pixel in the original field of view is subdivided into 2 × 2 subpixels. Orientations are shown with a color that corresponds to the computed orientation in an orientation color wheel. Pixels with a single orientation fill up all subpixels; pixels with two orientations fill the subpixels in a checkerboard pattern; pixels with three orientations populate the first three subpixels with an orientation, leaving the bottom right corner black. Additionally, the average scattering map (the mean along the *ϕ*_*c*_ direction) was used to determine the brightness of each pixel.

### 4.10 STA fiber orientation maps

The STA fiber orientation maps were computed from the high-resolution phase maps by performing a structure tensor analysis using the OrientationJ plugin for ImageJ [17] with a sigma-value of one pixel. The shown results in Figure 5 were plotted in HSV-space using the direction output as the hue and the energy of the structure tensor (its trace) as the value, leaving the saturation at one.

### 4.11 Fiber orientation difference maps

To compute the difference between the ComSLI fiber orientation maps from setup S1 and S2 (as shown in Figure 2), the lower-resolution maps from setup S2 were first upscaled with nearest-neighbor sampling to the same size as their S1 counterparts and then registered using the `imreg-dft` library [44]. The difference maps were computed by subtracting all combinations of the three direction maps obtained from the ComSLI analysis for both setups, then taking the combination with the smallest difference, and computing the average of the differences for all fiber orientations contained within one pixel. The difference maps show the absolute difference between the fiber orientations (from 0° to 90°) and are black in pixels where no orientation was found for setup S1 or S2; background is masked. In addition to the absolute difference maps, histograms of all angle difference values were plotted (from − 90° to 90°) for selected regions. For each histogram, the mean, mode, full width at half maximum (FWHM), and root mean square error (RMSE) were computed. The difference between ComSLI and STA fiber orientation (Figure 5) was computed in a similar way, by taking the smallest difference between all ComSLI fiber orientations and the STA orientation for each image pixel. The metrics (mean, mode, FWHM, RMSE) for the different histograms are shown in Supplementary Tables 1 and 2.

## AUTHOR CONTRIBUTIONS

Simon E. van Staalduine developed FP-SLM with input from Miriam Menzel, did the theoretical calculations and programming, performed the measurements with setup S2, applied FPM analysis to S2-images and ComSLI analysis to S1-images, and performed the STA analysis. Vittorio Bianco participated in the conception and design of the study, performed the measurements with setup S1, generated the high-resolution phase maps, and was involved in supervision. Pietro Ferraro acquired funding and was involved in supervision. Miriam Menzel participated in the conception and design of the study and was involved in supervision. Simon E. van Staalduine and Miriam Menzel wrote the first draft of the manuscript with inputs from Vittorio Bianco and Pietro Ferraro, and designed the figures. All authors revised the manuscript and approved the final version.

## ACKNOWLEDGMENTS

The authors thank Martin E. van Royen (Department of Pathology and Clinical Bioinformatics, Erasmus MC, Rotterdam, the Netherlands) and Wytske M. van Weerden (Department of Urology, Erasmus MC) for organizing the mouse samples, Corrina de Ridder and Debra Stuurman (Department of Urology, Erasmus MC) for tissue extraction, and Hamed Abbasi (Department of Otorhinolaryngology and Head and Neck Surgery, Erasmus MC) for the bright-field microscopy measurements. The tissue slide of the frog tadpole brain was prepared by Dott. Milena Kemali and available as part of the Archive Collection of the Istituto di Cibernetica, Neuroanatomy Laboratory, National Research Council, Italy. Further thanks go to Roger Woods from the UCLA Brain Research Institute for providing the vervet brain sample (National Institutes of Health under Grant Agreements no. R01MH092311 and 5P40OD010965), to the laboratory team at Forschungzentrum Jülich GmbH (INM-1), Germany, for preparing the vervet brain section, and to Bernd Rieger, Delft University of Technology, the Netherlands, for helpful comments.

## SUPPORTING INFORMATION

**Supplementary Figure 1:**
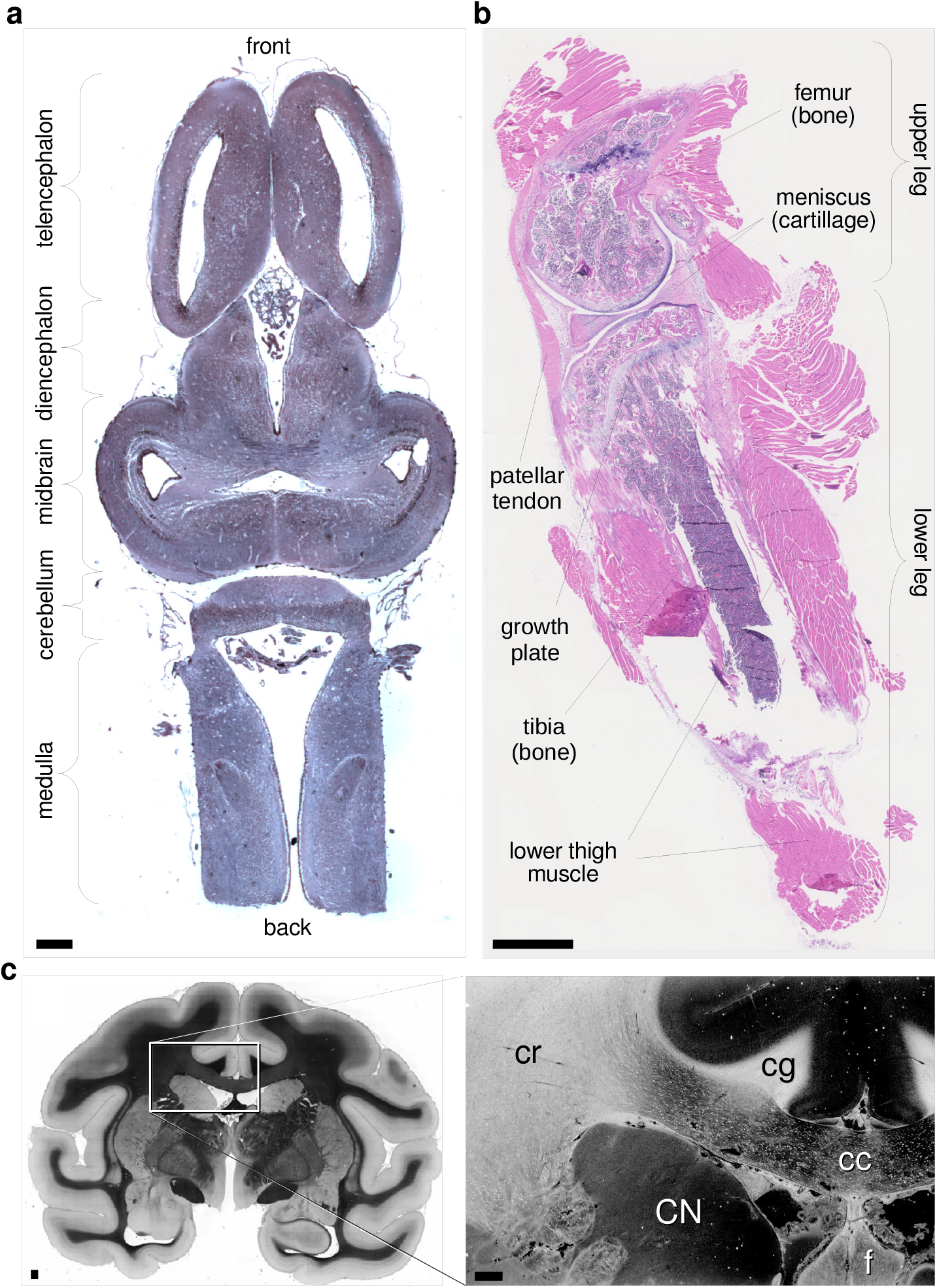
Measured samples with anatomical descriptions. a) Horizontal frog tadpole brain section. b) Longitudinal mouse knee section. c) Coronal vervet monkey brain section (bright-field image) with zoomed-in region (average scattering map), cr–corona radiata, cg–cingulum, cc–corpus callosum, f–fornix, CN–caudate nucleus. Scale bars are 1 mm.

**Supplementary Figure 2:**
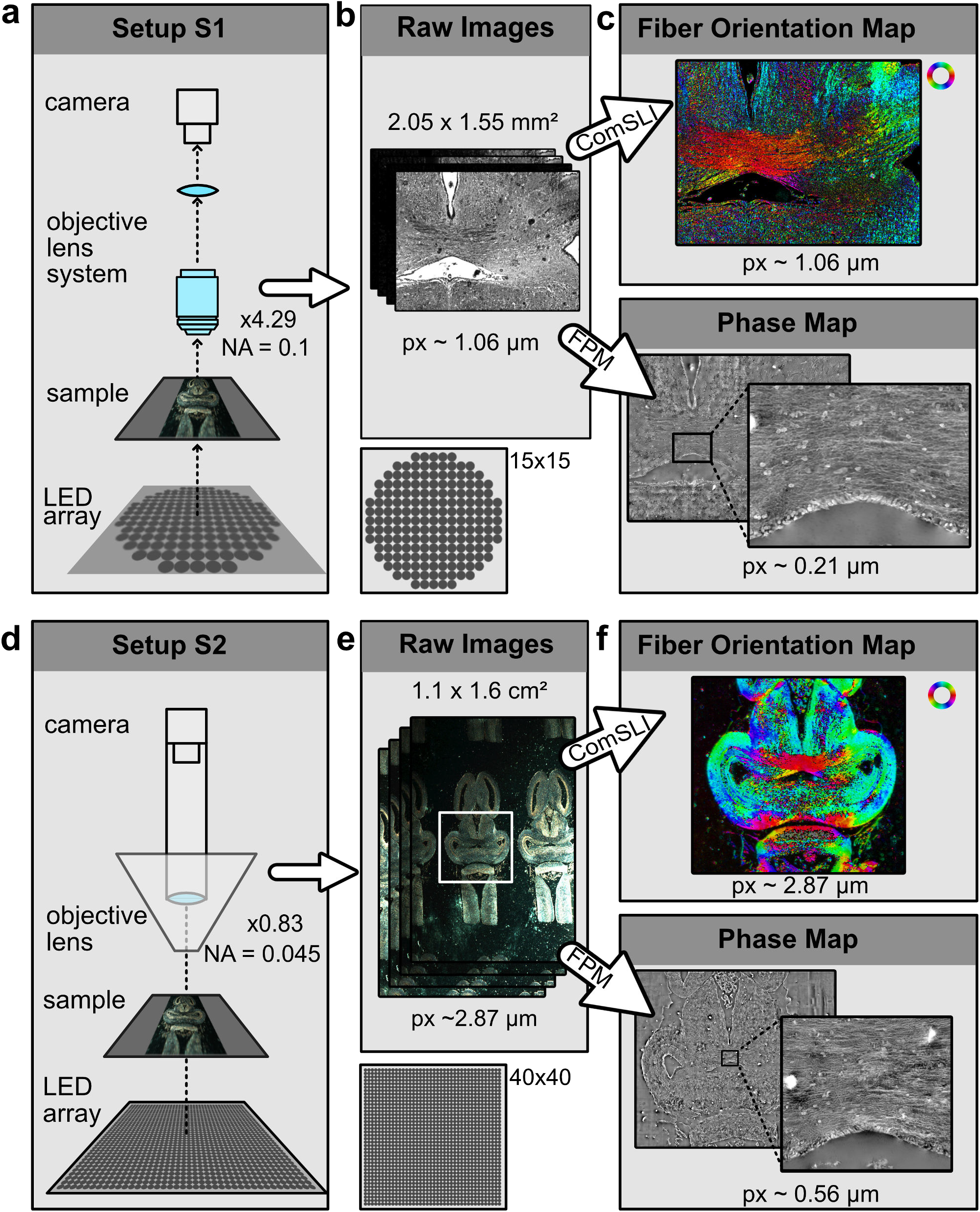
Comparison of the two setups used in this study: S1 (originally designed for FPM) and S2 (originally designed for ComSLI). a,d) Sketch of the two setups which employ different LED arrays and optical systems with different numerical apertures (NA) and magnifications. b,e) Image stacks resulting from a measurement with setup S1 and S2, with different fields of view (top) and pixel sizes (bottom). c,f) Corresponding fiber orientation maps (top) and phase maps (bottom) with different pixel sizes (px) obtained from a ComSLI and FPM analysis, respectively, shown for the marked regions.

**Supplementary Figure 3:**
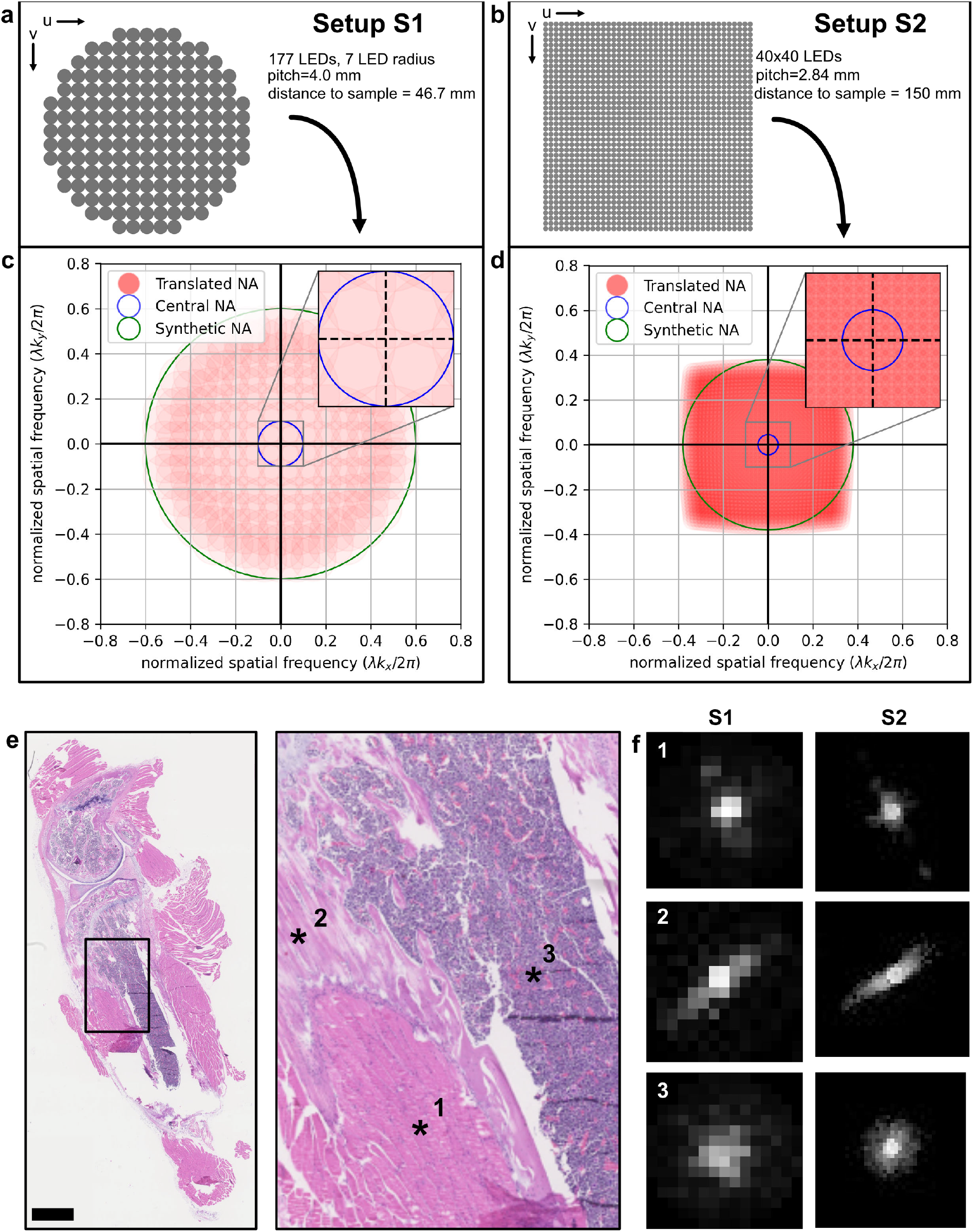
Scattering pattern generation. a),b) Layout of the LED arrays used in setup S1 (a) and in setup S2 (b). c),d) Frequency space coverage of the associated LED arrays (red), shown together with the coverage for direct illumination (blue) and synthetic numerical aperture (green). e) Zoom-in of the mouse knee histology section, showing three selected pixels: (1) muscle, (2) tendon, and (3) bone. Scale bar indicates 1 mm. f) Scattering patterns for the three selected pixels, obtained from setup S1 (left) and setup S2 (right). Tendon shows the highest degree of aligned structures (collagen fibers), resulting in a high-intensity line perpendicular to the fiber orientation (2). Bone is less intrinsically oriented and scatters more broadly than the other tissue types (3). Due to the different numbers of LEDs used for the measurements, S1 scattering patterns are more discretized than S2 scattering patterns, but they still correspond to each other.

**Supplementary Figure 4:**
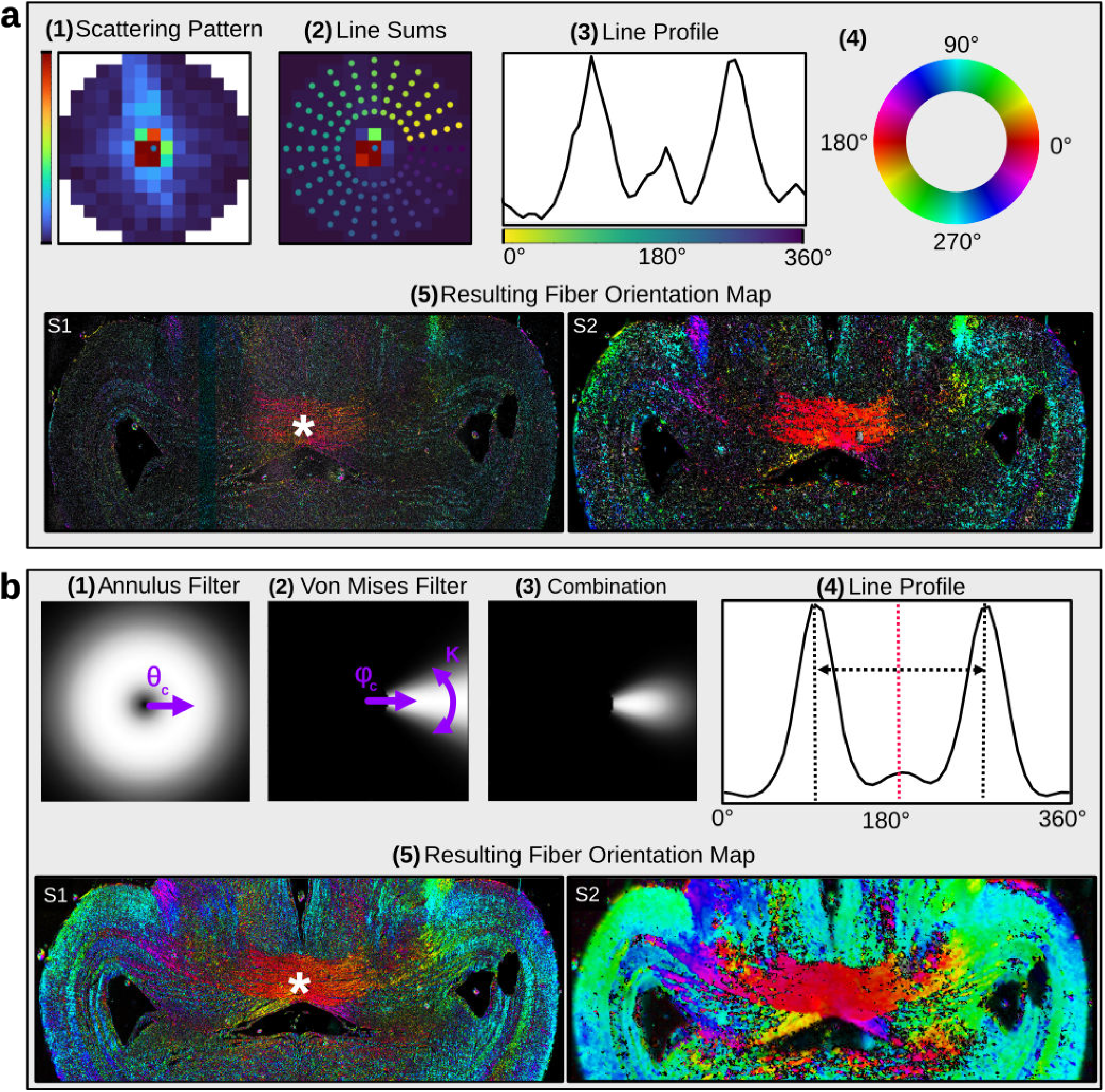
Computation of ComSLI fiber orientation maps without filtering (a) and with filtering (b) of scattering patterns. a) Without filtering, each scattering pattern (1) is sampled from an inner to an outer circle along a line in 15° azimuthal steps as shown in (2), and for each azimuthal angle the sum of intensities is plotted (3). The fiber orientation is then computed from the mid-position of a peak pair in the line profile and represented by a color according to the color wheel (4). This procedure is performed for each pixel, resulting in a fiber orientation map (5), which is shown here exemplary for the mid region of the frog brain section for setup S1 and S2. b) With filtering, an annulus filter (1) and von Mises filter (2) are combined (3) and applied to the scattering pattern for the different azimuthal angles, resulting in a smoothed line profile (4). In this profile, it is much easier to reliably determine the position of prominent peaks (black dashed lines) and their mid-position (red dashed line) corresponding to the fiber orientation in the pixel, here a horizontal orientation encoded in red in the fiber orientation map (5), marked by the asterisk.

**Supplementary Table 1:**
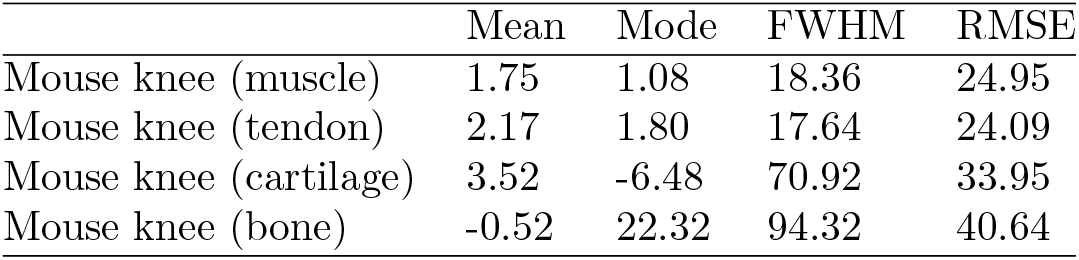
Histogram metrics of main Figure 2 (in degrees), showing the difference between ComSLI fiber orientations obtained from setup S1 and setup S2: mean, mode, full width at half maximum (FWHM), and root mean square error (RMSE).

**Supplementary Table 2:** Histogram metrics of main Figure 5 (in degrees), showing the difference between ComSLI and STA fiber orientations for setup S1 and S2: mean, mode, full width at half maximum (FWHM), and root mean square error (RMSE).

|  | Mean | Mode | FWHM | RMSE |
| --- | --- | --- | --- | --- |
| Frog brain (S1) | 0.28 | -2.88 | 79.56 | 40.48 |
| Frog brain (S2) | 5.90 | -2.16 | 60.84 | 25.80 |
| Mouse knee (S1) | -6.21 | -8.64 | 24.48 | 36.38 |
| Mouse knee (S2) | 0.99 | -3.24 | 30.96 | 32.43 |
| Vervet brain (S2) | -5.82 | -6.12 | 53.64 | 27.11 |

## References

[1] H. Blom, J. Widengren, Chemical Reviews 2017, 117, 11 7377, publisher: American Chemical Society.

[2] G. Lukinavičius, J. Alvelid, R. Gerasimaitė, C. Rodilla-Ramirez, V. T. Nguyn, G. Vicidomini, F. Bottanelli, K. Y. Han, I. Testa, Nature Reviews Methods Primers 2024, 4, 1 1, publisher: Nature Publishing Group.

[3] A. D. Elliott, Current Protocols in Cytometry 2020, 92, 1 e68.

[4] R. K. Benninger, D. W. Piston, Current Protocols in Cell Biology 2013, 59, 1 4.11.1, eprint: https://currentprotocols.onlinelibrary.wiley.com/doi/pdf/10.1002/0471143030.cb0411s59.

[5] G. Zheng, R. Horstmeyer, C. Yang, Nature Photonics 2013, 7, 9 739.

[6] S. Jian, K. Guo, J. Liao, G. Zhen, Biomedical Optics Express 2018, 9 3306.

[7] S. Xu, X. Yang, P. Ritter, X. Dai, K. C. Lee, L. Kreiss, K. C. Zhou, K. Kim, A. Chaware, J. Neff, C. Glass, S. A. Lee, O. Friedrich, R. Horstmeyer, Advanced Photonics 2024, 6, 2 026004.

[8] Y. Rivenson, T. Liu, Z. Wei, Y. Zhang, K. de Haan, A. Ozcan, Light: Science & Applications 2019, 8, 1 23, publisher: Nature Publishing Group.

[9] N. Borhani, A. J. Bower, S. A. Boppart, D. Psaltis, Biomedical Optics Express 2019, 10, 3 1339.

[10] Y. Rivenson, H. Wang, Z. Wei, K. de Haan, Y. Zhang, Y. Wu, H. Gönaydin, J. E. Zuckerman, T. Chong, A. E. Sisk, L. M. Westbrook, W. D. Wallace, A. Ozcan, Nature Biomedical Engineering 2019, 3, 6 466, publisher: Nature Publishing Group.

[11] M. E. Kandel, Y. R. He, Y. J. Lee, T. H.-Y. Chen, K. M. Sullivan, O. Aydin, M. T. A. Saif, H. Kong, N. Sobh, G. Popescu, Nature Communications 2020, 11, 1 6256, publisher: Nature Publishing Group.

[12] V. Bianco, M. D’Agostino, D. Pirone, G. Giugliano, N. Mosca, M. Di Summa, G. Scerra, P. Memmolo, L. Miccio, T. Russo, E. Stella, P. Ferraro, Small Methods 2023, 7, 11 2300447, eprint: https://onlinelibrary.wiley.com/doi/pdf/10.1002/smtd.202300447.

[13] Y. Shi, A. Toga, Molecular Psychiatry 2017, 22 1230.

[14] J. N. Ouellette, C. R. Drifka, K. B. Pointer, Y. Liu, T. J. Lieberthal, W. J. Kao, J. S. Kuo, A. G. Loeffler, K. W. Eliceiri, Bioengineering 2021, 8, 2.

[15] H. Su, M. Karin, Trends in Cancer 2023, 9, 9 764.

[16] W. Han, S. Chen, W. Yuan, Q. Fan, J. Tian, X. Wang, L. Chen, X. Zhang, W. Wei, R. Liu, J. Qu, Y. Jiao, R. Austin, L. Liu, PNAS 2016, 113, 40 11208.

[17] Z. Püspöki, M. Storath, D. Sage, M. Unser, In W.H. De Vos, S. Munck, J.-P. Timmermans, editors, Focus on Bio-Image Informatics, volume 219, 69–93. Springer International Publishing, Cham, ISBN 978-3-319-28547-4 978-3-319-28549-8, 2016.

[18] M. Axer, K. Amunts, D. Grässel, C. Palm, J. Dammers, H. Axer, U. Pietrzyk, K. Zilles, NeuroImage 2011, 54, 2 1091.

[19] M. Georgiadis, F. a. d. Heiden, H. Abbasi, L. Ettema, J. Nirschl, H. M. Taghavi, M. Wakatsuki, A. Liu, W. H. D. Ho, M. Carlson, M. Doukas, S. A. Koppes, S. Keereweer, R. A. Sobel, K. Setsompop, C. Liao, K. Amunts, M. Axer, M. Zeineh, M. Menzel, Micron-resolution fiber mapping in histology independent of sample preparation, 2024, URL https://www.biorxiv.org/content/10.1101/2024.03.26.586745, Pages: 2024.03.26.586745 Section: New Results.

[20] M. Dohmen, M. Menzel, H. Wiese, J. Reckfort, F. Hanke, U. Pietrzyk, K. Zilles, K. Amunts, M. Axer, NeuroImage 2015, 111 464.

[21] M. Menzel, M. Axer, H. De Raedt, I. Costantini, L. Silvestri, F. S. Pavone, K. Amunts, K. Michielsen, Physical Review X 2020, 10, 2 021002, publisher: American Physical Society.

[22] M. Menzel, J. A. Reuter, D. Gräßel, M. Huwer, P. Schlömer, K. Amunts, M. Axer, NeuroImage 2021, 233 117952.

[23] M. Menzel, M. Ritzkowski, J. A. Reuter, D. Gräßel, K. Amunts, M. Axer, Frontiers in Neuroanatomy 2021, 15.

[24] V. Bianco, L. Miccio, D. Pirone, E. Cavalletti, J. Behal, P. Memmolo, A. Sardo, P. Ferraro, Scientific Reports 2024, 14, 8418.

[25] V. Bianco, B. Mandracchia, J. Bėhal, D. Barone, P. Memmolo, P. Ferraro, IEEE Journal of Selected Topics in Quantum Electronics 2021, 27, 4 1, conference Name: IEEE Journal of Selected Topics in Quantum Electronics.

[26] M. Menzel, D. Gräßel, I. Rajkovic, M. M. Zeineh, M. Georgiadis, eLife 2023, 12 e84024, publisher: eLife Sciences Publications, Ltd.

[27] T. Aidukas, R. Eckert, A. R. Harvey, L. Waller, P. C. Konda, Scientific Reports 2019, 9, 1 7457, publisher: Nature Publishing Group.

[28] R. Horstmeyer, J. Chung, X. Ou, G. Zheng, C. Yang, Optica 2016, 3, 8 827–.

[29] R. Schurr, A. A. Mezer, Science 2021, 374, 6568 762, publisher: American Association for the Advancement of Science.

[30] C. Sampias, H&E Staining Basics: Troubleshooting Common H&E Stain Problems, 2024, URL https://www.leicabiosystems.com/en-nl/knowledge-pathway/he-basics-part-4-troubleshooting-he/.

[31] M. Kemali, V. Braitenberg, Atlas of the Frog’s Brain, Springer, 1970.

[32] M. Valentino, D. Pirone, J. Béhal, M. Mugnano, R. Castaldo, G. C. Lama, P. Memmolo, L. Miccio, V. Bianco, S. Grilli, P. Ferraro, Journal of Physics: Photonics 2024, 6, 1 015004.

[33] G. Zheng, C. Shen, S. Jiang, P. Song, C. Yang, Nature Reviews Physics 2021, 3, 3 207.

[34] L. Tian, Z. Liu, L.-H. Yeh, M. Chen, J. Zhong, L. Waller, Optica 2015, 2, 10 904.

[35] L. Loetgering, T. Aidukas, K. C. Zhou, F. Wechsler, R. Horstmeyer, Microscopy Today 2022, 30, 5 36.

[36] J. Arines, Applied Optics 2003, 42, 17 3373.

[37] D. Pirone, V. Bianco, M. Valentino, M. Mugnano, V. Pagliarulo, P. Memmolo, L. Miccio, P. Ferraro, Optics and Lasers in Engineering 2022, 156 107103.

[38] L. Loetgering, M. Du, D. B. Flaes, T. Aidukas, F. Wechsler, D. S. P. Molina, M. Rose, A. Pelekanidis, W. Eschen, J. Hess, T. Wilhein, R. Heintzmann, J. Rothhardt, S. Witte, Optics Express 2023, 31, 9 13763, publisher: Optica Publishing Group.

[39] T. Aidukas, L. Loetgering, A. R. Harvey, Optics Express 2022, 30, 13 22421, publisher: Optica Publishing Group.

[40] S. Preibisch, S. Saalfeld, P. Tomancak, Bioinformatics 2009, 25, 11 1463.

[41] M. Menzel, S. F. Pereira, Biomedical Optics Express 2020, 11, 8 4735, publisher: Optica Publishing Group.

[42] M. Kittisopikul, A. Vahabikashi, T. Shimi, R. D. Goldman, K. Jaqaman, Bioinformatics 2020, 36, 20 5093.

[43] J. A. Reuter, M. Menzel, Journal of Open Source Software 2020, 5, 54 2675.

[44] B. Reddy, B. Chatterji, IEEE Transactions on Image Processing 1996, 5, 8 1266, conference Name: IEEE Transactions on Image Processing.

